# Profiling of Aggregated Amyloid β42-induced Proteomic Alterations in KOLF2.1J Induced Pluripotent Stem Cell-derived Neurons

**DOI:** 10.64898/2026.09.20.753014

**Authors:** Benjamin Jin, Ziyi Li, Ying Hao, Isabelle Kowal, Jacob Epstein, Marianita Santiana, Cory A. Weller, Mike A. Nalls, Dingyin Tao, Shengyun Fang, Priyanka Narayan, Andrew B. Singleton, Kendall Van Keuren-Jensen, Luigi Ferrucci, Michael E. Ward, Mark R. Cookson, Erika Lara, Veronica H. Ryan, Yue Andy Qi

## Abstract

Amyloid β (Aβ) plaques are a hallmark of Alzheimer’s disease (AD). A human cellular neuronal model that recaptures Aβ-induced pathology is critical for advancing AD research. However, comprehensive proteomic profiling of Aβ-induced cellular model remains elusive. In this study, we investigated the proteomic changes in induced pluripotent stem cell (iPSC)-derived neurons (iNs) exposed to synthetic Aβ (1-42) peptides (Aβ42A) to improve our understanding of the cellular responses of Aβ aggregates and to establish a human-related platform for AD research. Aβ42A formed extracellular aggregates around neuronal soma and neurites, impaired neurite outgrowth, and induced the expression of multiple AD-associated genes, such as APOE, BACE1, ADAM10. To define the molecular landscape of Aβ-induced neuronal dysfunction, we performed proteomic analyses of whole-cell lysates as well as soma- and neurite-enriched fractions. Proteomic profiling revealed extensive gene alterations in pathways associated with synaptic function, neuronal maintenance, and AD pathogenesis, such as the upregulation of APOE. Importantly, the Aβ42A-iNs system recapitulated molecular signatures observed in AD brain tissue and cerebrospinal fluid. Representative protein changes, including APOE upregulation and its colocalization with Aβ aggregates, were further validated in postmortem human AD brain tissue. Together, these findings manifest that the Aβ42A-iNs system reproduces multiple cellular and molecular features of AD and exhibits strongly consensus with clinical observations. It provides a valuable platform for investigating Aβ-driven neurodegeneration and for the discovery of therapeutic targets for AD.

## Background

The increase in the incidences of Alzheimer’s disease (AD) globally and the lack of effective treatments for AD urge the imminent need for new therapeutics with the improved understanding of the pathogenesis of AD. Extracellular amyloid β (Aβ) plaques have long been a pathogenetic and therapeutic target for AD [1]. The primary component of Aβ plaques is Aβ 1-42 peptides (Aβ42), which aggregate into oligomers, fibrils, and insoluble plaques, leading to neuronal dysfunction [2]. Various strategies have been employed to generate neurotoxic Aβ aggregates *in vitro* using Aβ42 peptides with limited success [3]. Human induced pluripotent stem cells (iPSCs) are powerful tools for experimental disease modeling in translational research, including studies of neurodegenerative diseases [4, 5]. The successful establishment of the iPSC-derived neuron differentiation culture method has significantly promoted the study of neurodegenerative diseases, such as AD [5, 6].

Synthetic Aβ42 peptides and iPSC-derived neurons (iNs) have been used to advance our understanding of the pathophysiology of AD [7–9]. Although a cellular system with Aβ-induced neurotoxicity has been reported to study AD pathology and identify potential therapeutic targets, this cellular system has not been well-characterized at the proteomic level, thus reducing its translational application [10]. Therefore, we established a platform incorporating synthetic Aβ42 and iNs derived from a well-characterized iPSC line, KOFL2.1J, to recapitulate the neuronal pathophysiology of AD induced by Aβ aggregates in vitro. Through cellular imaging analyses and proteomic profiling, we demonstrated that our Aβ42A-iNs system reproduced key AD-related pathophysiological features, including Aβ aggregates, neurite fragmentation, downregulation of neuronal markers, and upregulation of multiple AD-associated risk factors. We identified common regulatory proteins across whole neurons, soma, and neurites in our system. Furthermore, several of these molecules are also detected in postmortem brain tissue and cerebrospinal fluid from AD patients. Overall, our platform provides a spatially resolved proteomic profile of a cellular AD model and reveals novel potential therapeutic targets, such as PTN and MDK.

## Materials and Methods

### iPSC-derived neuron culture and Aβ treatment

KOLF2.1J iPSCs were transfected with human neurogenin-2 (NGN2) vectors, differentiated into neurons, and maintained as previously described [11, 12]. Briefly, KOLF2.1J iPSCs were transfected with NGN2 vectors and selected with puromycin. KOLF2.1J_NGN2 transfectants were cultured in Matrigel-coated 6-well plates with DMEM/F12 media supplemented with N2 (1:100), Non-Essential Amino Acid (1:100), GlutaMAX (1:100), and doxycycline (2 µg/mL) to start differentiation. On day 4, the immature neurons were lifted with Accutase and seeded in poly-L-ornithine (PLO)-coated plates with neuronal maturation media (NMM) containing DMEM/F12:BrainPhys Neuronal Media (1:1) supplemented with N21 MAX (1:50), GDNF (10 ng/mL), BDNF (10 ng/mL), NT-3 (10 ng/mL), laminin (1 µg/mL), doxycycline (2 µg/mL), uridine (1 µM), and 5-fluoro-2’-deoxyuridine (1 µM). Starting on day 7, half of the media was replaced every 3-4 days with NMM containing BrainPhys Neuronal Media supplemented with all required additives. Lyophilized AggreSure Aβ1-42 peptides (which was called Aβ42A in the manuscript. Anaspec: AS-72216) were resuspended in DMSO and then diluted with phosphate-buffered saline (PBS) to form a 100 µM solution. The iNs were treated with Aβ42A at the concentrations and durations detailed in the figure legends and then harvested for various analytic assays.

### Neurite outgrowth assay

KOLF2.1J iPSCs with NGN2 were co-transduced with lentiviral cytoplasmic mScarlet (MK-EF1a-mScarlet) and nuclear mNeonGreen (MK-EF1a-mNeonGreen-NLS) and differentiated into iNs as previously described [12, 13]. Briefly, unlabeled KOLF2.1J_NGN2 iPSCs and fluorescent KOLF2.1J_NGN2 iPSCs were differentiated into immature neurons in Matrigel-coated plates, respectively. On day 4, these two immature neurons were lifted with Accutase to generate single cell suspensions. A mixture of 98-99% unlabeled neurons and 1-2% fluorescent neurons was seeded in PLO-coated plates for maturation and were treated with Aβ42A. The course of the neuronal differentiation and neurite outgrowth was monitored with an Incucyte S3 Live-Cell Analysis System using a 10X or 20X objective every 24 h at 37°C; phase and fluorescence images were acquired at each time point. The NeuroTrack Incucyte software module was used to analyze neurite outgrowth.

### Western blot

Aβ peptides (10 ng) were separated on 4-12% SDS-PAGE (Bio-Rad: 4561094) and transferred to nitrocellulose membrane using Bio-Rad Trans-Blot semi-dry transfer system. The transferred membrane was blocked with a Tris-buffered saline (TBS) solution containing 5% BSA and 0.2% v/v Tween-20 and blotted with 1:1000 mouse 6E10 primary antibody (Biolegend) overnight at 4 °C following 1:3000 HRP-conjugated Goat-anti-mouse secondary antibody (Thermo Fisher: G-21040). The membrane was incubated with SuperSignal™ Substrate (Thermo Fisher: A38556) and imaged using a ChemiDoc MP Imaging System (Bio-Rad).

### Immunocytochemistry and confocal imaging

The iNs were fixed with 4% PFA (Thermo Fisher: 15710), permeabilized with 0.1% Triton-X 100, and blocked in 2% bovine serum albumin (BSA, Jackson ImmunoResearch 001000173). The samples were incubated with primary antibodies at 4 °C overnight, followed by incubation with fluorescence-labeled secondary antibodies incubation at room temperature for one hour. Images were acquired with a Nikon Ti2 confocal microscope with a 60X water immersion objective, following the manufacturer’s standard imaging protocol. The images were processed with Imaris software (Oxford Instruments). The primary antibodies were anti-Aβ antibody (1:100 dilution (10 μg/mL), Biolegend: 803001, 6E10) and anti-MAP2 antibody (1:500 dilution, Thermo Fisher: PA1-10005). The secondary antibodies were goat-anti-mouse Alexa Fluor 405 and goat-anti-chicken Alexa Fluor 405 and used at a dilution of 1:500.

Human brain tissue sections were purchased from Biochain. The sections were fixed and stained with primary antibodies and secondary antibodies following the standard protocol. The primary antibodies are anti-Aβ antibody (1:50 dilution, CST: 8243) and anti-APOE (1:20 dilution, CST: 74417). The secondary antibodies were goat-anti-rabbit Alexa Fluor 488 and goat-anti-mouse Alexa Fluor 647 and used at a dilution of 1:500.

Images were acquired with a Nikon Ti2 confocal microscope with a 20X objective, following the manufacturer’s standard imaging protocol. The images were processed with Imaris software.

### Proteomic sample preparation

Sample preparation for proteomics was conducted on a fully automated workflow as previously reported [14, 15]. Briefly, iNs samples (n=6) were washed with PBS and a 200 μL of SP3 lysis buffer containing proteasome inhibitors was added. The lysates were incubated at 65°C for 30 minutes allowing for protein denaturation and followed by alkylation with 10 mM iodoacetamide in the dark. Protein enrichment and on-bead tryptic digestion were conducted via a KingFisher APEX robotic system. Proteins were digested with a trypsin/Lys-C mixture in 50 mM ammonium bicarbonate with a ratio of 1:20 (trypsin: protein) at 37°C for 16 hours. After digestion, the peptides were vacuum-dried and reconstituted in 2% acetonitrile (ACN) with 0.1% formic acid. Approximately 1 μg of digested peptide was used for proteomic analysis.

### Liquid chromatography and mass spectrometry (LC/MS)

The digested peptides were analyzed with an UltiMate 3000 nano-HPLC system (Thermo Fisher) coupled with an Orbitrap Eclipse mass spectrometer (Thermo Fisher). Peptide separation was performed on an ES903A nanocolumn (Thermo Fisher 75 μm × 500 mm, 2 μm C18 particle size) using an 80-minute linear gradient of 2% - 40% phase A (0.1% formic acid in water) and phase B (5% DMSO in 0.1% formic acid, in ACN). The column temperature was maintained at 60°C, and the flow rate was 300 nL/min. Data were acquired in data-independent acquisition (DIA) mode. The MS1 scan was set at a resolution of 120,000, with a standard AGC target, and the maximum injection time was set to auto. The MS2 scans covered a precursor mass range of 400–1000 m/z, using an isolation window of 8 m/z with 1 m/z overlap, resulting in a total of 75 windows per scan cycle. Fragmentation was performed via high-energy collisional dissociation (HCD) with a normalized collision energy of 30%. MS2 spectra were acquired at a resolution of 30,000, with a scan range of 145–1450 m/z, and the loop control was set to 3 seconds.

### Neuritic fractions (NFs) and soma-enriched fractions (SEFs) proteomics

KOLF2.1J iPSC-derived neurons were cultured in a Boyden chamber as previously described [16]. Briefly, the Boyden chamber insert was coated with PLO and dried; the bottom of the insert was coated with 10 μg/mL laminin prior to the iNs were plated. On day 4, the iNs were plated, and half of the neuron culture media were changed from both the top and bottom of the Boyden chamber every 3 days for one week. On Day 15, the iNs were treated with 2.5 μM Aβ42A for 7 days. Half of the media were changed every 3 days for both treated and untreated wells. On Day 21, the iNs were washed with PBS, and neuritic fractions (NFs) were first collected by scraping neurites from the bottom of the insert, followed by soma-enriched fractions (SEFs) collection from the top of the insert. The NFs and SEFs were placed in separate tubes and pelleted by centrifugation, after which the PBS was removed. The NFs and SEFs were flash-frozen before proceeding to mass spectrometry preparation.

### Proteomics database search

We conducted a DIA database search via Spectronaut (version 19, Biognosys) with the directDIA search strategy. We used the reviewed UniProt human proteome reference FASTA file with one protein per gene (20,384 entries). For proteomic analysis, we applied the factory default settings. Trypsin/P and Lys-C were selected as the digestion enzymes. The digestion specificity was defined as specific, permitting a maximum of two missed cleavages per peptide. Carbamidomethylation of cysteine residues was set as a fixed modification. For variable modifications, oxidation of methionine and N-terminal acetylation were considered. The false discovery rate (FDR) thresholds for peptide-spectrum matches (PSMs), peptide, and protein groups were set to 0.01. Cross-run normalization was enabled to account for systematic variations across runs and no imputation was performed on the dataset. Unless explicitly stated, all other parameters were maintained as default Spectronaut settings.

### Data visualization and statistical analysis

Further downstream analyses were performed using our in-house proteomics analysis pipeline, ProtPipe [17]. Proteins were considered differentially expressed if the absolute value of the log2 fold change was greater than 0.5 and the adjusted *p*-value was less than 0.05, with multiple testing correction applied via the Benjamini-Hochberg method. Functional enrichment analysis was performed via the *clusterProfiler* R package within the ProtPipe workflow, with pathways considered significant at a q-value threshold of less than 0.05.

To investigate the translational relevance of our Aβ42A-iNs system, we searched and found a publication that conducted a comprehensive meta-analysis of proteomics data from postmortem brain tissues of AD patients across 7 datasets [18]. We identified overlapping differentially expressed proteins (DEPs) in our findings and those from AD patients using a cutoff of absolute log2 fold change greater than 0.5 and an adjusted *p*-value less than 0.05. All visualizations were generated via the ggplot2 and pheatmap packages in R (version 4.3.1).

## Results

### Aβ42A aggregate-induced neurotoxicity in KOLF2.1J iPSC-derived neurons

Aβ aggregates are one of AD pathogenetic features and can induce neurotoxicity in cultured neurons in vitro [19, 20]. To validate that our Aβ42A consists of aggregates, we run Western blot with Aβ42A and probed with an anti-Aβ antibody. Aβ42A exhibited multiple bands above 37 kDa, indicating the presence of aggregates in Aβ42A (Fig .1 A). Extracellular Aβ aggregates can cause neurotoxicity in various ways, such as the loss of neuronal markers, dendrite reduction, and axon fragmentation [20]. To validate that Aβ42A can cause neurotoxicity, we treated iNs with Aβ42A in iNs transfected with cytosolic fluorescence protein (mScarlet) and assessed neurite outgrowth. Live imaging demonstrated that Aβ42A progressively impaired neurite outgrowth (Aβ42A vs VC, *p*=0.0049, n=6). (Fig. 1C & D). These observations demonstrated that we successfully generated neurotoxic Aβ aggregates. To confirm that Aβ42A formed aggregates in the *in vitro* culture system, Aβ42A-treated iNs were stained with anti-Aβ and anti-MAP2 antibodies. A variety of different sizes of clusters indicating Aβ42 aggregates were observed (Fig. 1E II) and accumulated extracellularly (Fig. 1E V). Aβ42A-induced neurotoxicity were shown as the reduced expression of MAP2 (Fig. 1E IV).

**Fig. 1.**
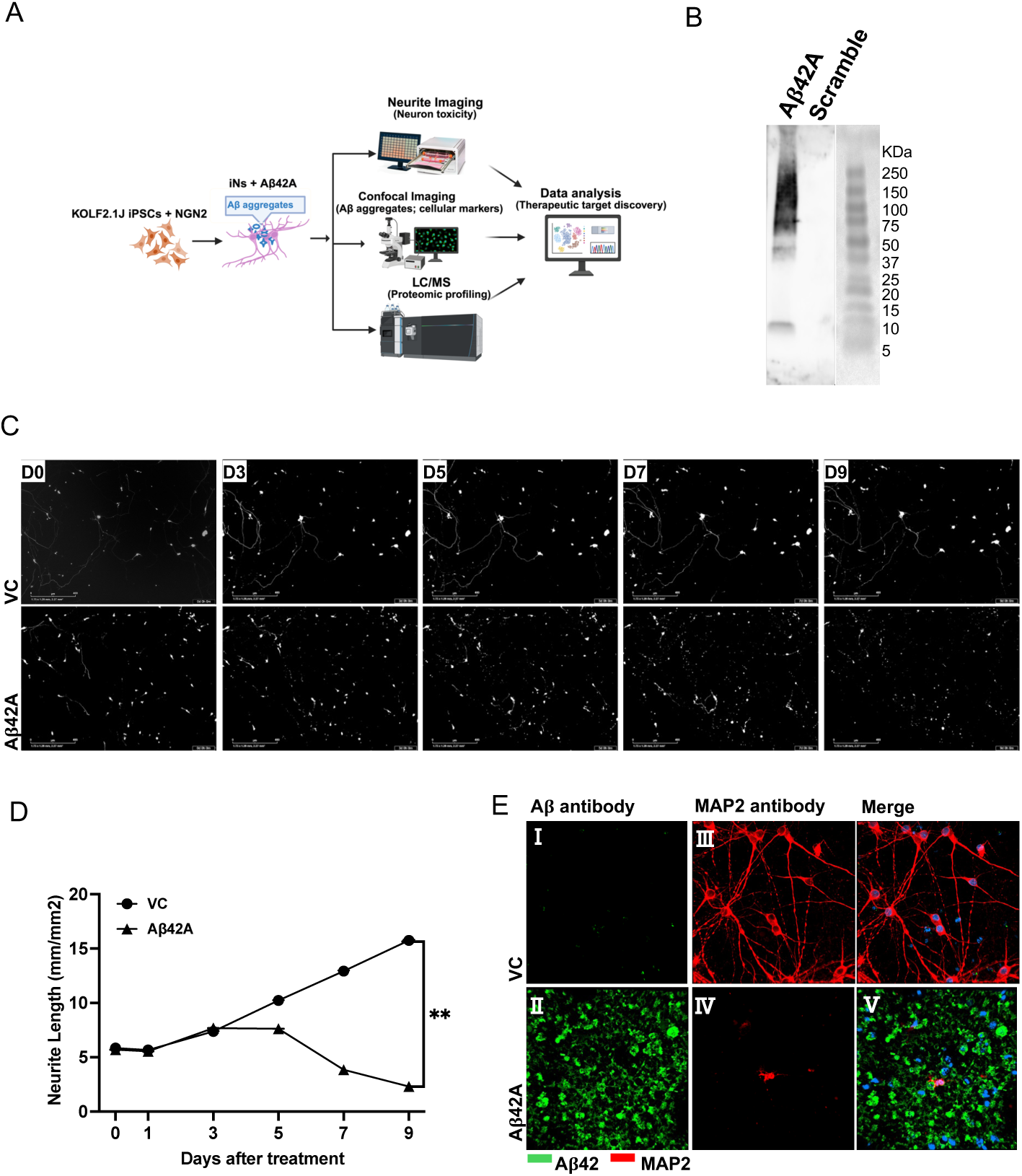
Aβ42A elicited neurotoxicity in KOLF2.1J iPSC-derived neurons. A. Scheme of the experimental design. KOLF2.1J iPSCs were transfected with NGN2 and differentiated into neurons according to the standard protocol developed by the Center for Alzheimer’s and Related Dementias. The iNs were treated with Aβ42 peptides and assayed via confocal microscopy imaging and real-time live imaging technologies, and mass spectrometry. The data were processed and analyzed with a range of platforms and computational methodologies. B. Western blot of Aβ42 peptides. Aβ42A and scramble peptides were resolved by SDS-PAGE and transferred to nitrocellulose membrane for immunoblot analysis. The peptide-bound membrane was probed with anti-Aβ antibody (6E10) firstly and then incubated with horseradish peroxidase (HRP)-conjugated secondary antibodies. The developed membrane was imaged using a ChemiDoc imaging system. C. Aβ42A disrupted neurite outgrowth. iNs expressing cytosolic mScarlet and nuclear mNeonGreen were treated with Aβ42A at a dose of 5 µM. The neurite outgrowth was monitored by the IncuCyte system. A representative image showed neurite outgrowth across the specific time points. The neurites continued growing in the vehicle (VC)-treated groups, whereas the outgrowth of neurites was disrupted in the Aβ42A-treated iNs. D. The length of outgrowth neurites. Aβ42A treatment resulted in a significant reduction in neurite length compared to controls (**, *p*=0.0049) (n=6). E. Aβ42A formed aggregates and reduced MAP2 expression in the iNs. Day 7 iNs were treated with 5 µM Aβ42A for 7 days. The samples were fixed and stained with anti-Aβ and anti-MAP2 antibodies and followed by incubation with Alexa 488- and Alexa561-conjugated secondary antibodies. The images were taken with a Nikon Ti2 confocal microscope with a 60x objective. Aβ42A formed aggregates in the Aβ42A-treated samples (II) but not in the VC-treated samples (I). Aβ42A treatment induced neurotoxicity with the reduced MAP2 expression (IV) and Aβ42A aggregates were deposited extracellularly (V).

### Replication of characteristic modulations of AD-associated risk genes and cellular responses

Mass spectrometry-based proteomics plays an increasing role in the identification of biomarkers and therapeutic targets and the understanding of disease mechanisms in AD [18, 21]. The complexity of AD pathogenesis implies the involvement of multiple genes and cellular pathways [22]. Reports have shown dysregulation of neuronal markers, such as MAP2, TUBB3, NEF, and SYN, indicating neurodegeneration [23–26]. Additionally, certain genes, including APOE, BACE1, and ADAM10, have been recognized as risk factors for AD [27, 28]. To investigate the cellular responses of KOLF2.1J iNs to Aβ42A, we used mass spectrometry-based proteomics to study changes in cellular components and pathways following Aβ42A treatment. We found that Aβ42A-treated iNs displayed neuronal changes characteristic of AD pathologies. Neuronal damage was manifested as a reduction in MAP2, TUBB3, NEFH, NEFM, NEFL, and SYN3 expression (Fig. 2A). AD-associated risk genes, including APP, APOE, ADAM10, BACE1, PSEN2, and CLU, were significantly increased (Fig. 2B). Differential protein abundance analysis further revealed more genes that may be involved in the neurotoxic processes induced by Aβ42A (Fig. 2C), including the increased SFRP1, SPOCK2, FBLN1, PTN, and APOA1, et al, as well as the decreased TLL2, PNN, ARL6P4, RAMAC, and NTM, et al.

**Fig. 2.**
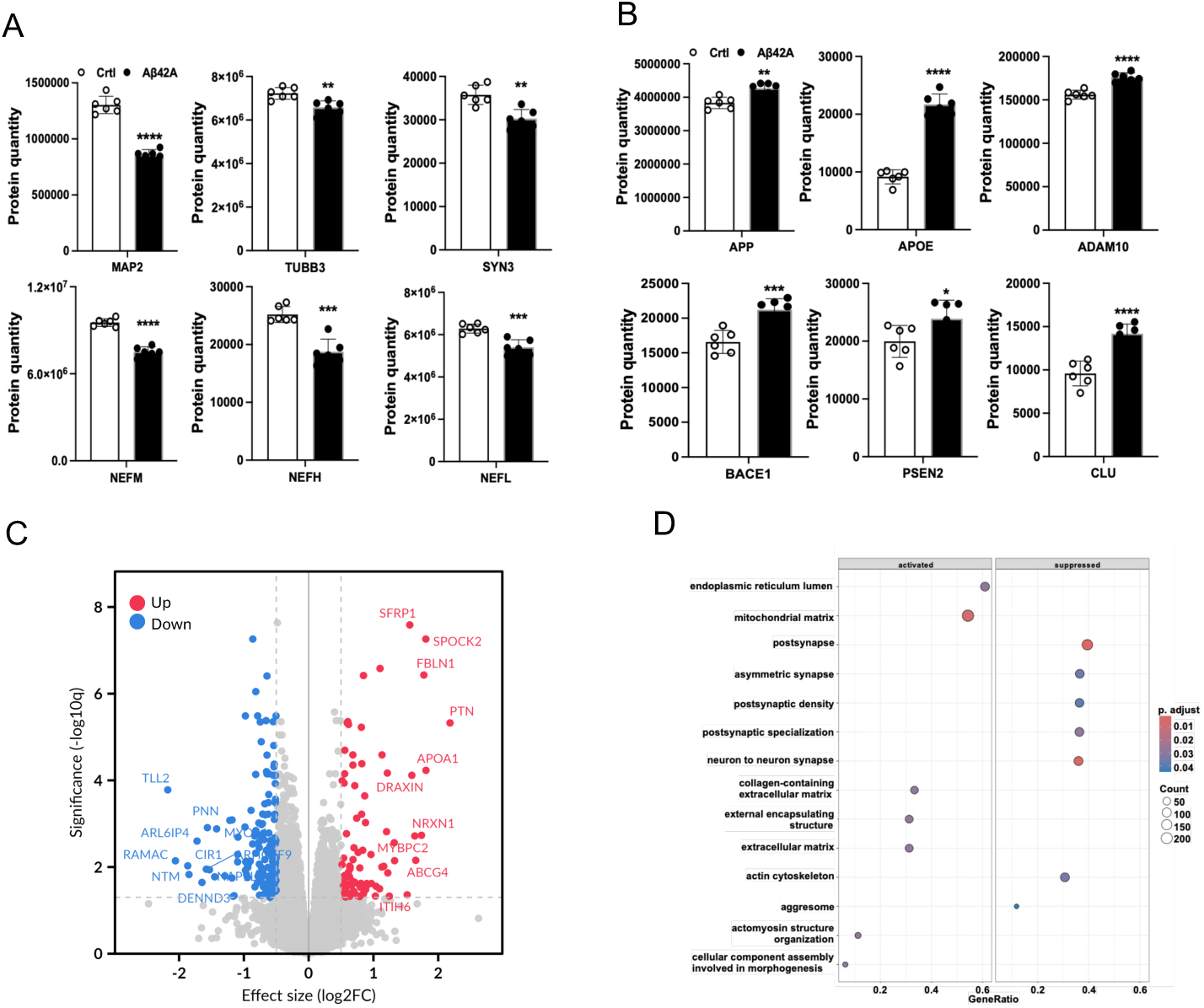
Aβ42A induced pathophysiological responses of AD in KOLF2.1J iPSC-derived neurons. A. Aβ42A modulated the expression of multiple AD-related genes and cellular pathways in KOLF2.1J iNs. Day 7 iNs were treated with 5 µM Aβ42A for 7 days. The iNs were collected for LC-MS assay and proteomics analysis. Aβ42A treatment significantly downregulated multiple mature neuron markers, including MAP2, TUBB3, SYN3, NEFH, NEFM, and NEFL. B. Aβ42A treatment significantly upregulated several AD-associated molecules, including APP, APOE, ADAM10, BACE1, PSEN2 and CLU. C. Volcano plot of differential protein abundance analysis indicating the upregulated and downregulated proteins. D. Thematic classification of GO terms affected by Aβ42A treatment.

To identify the cellular pathways involved in Aβ42A-induced neurotoxicity, we performed Gene Ontology (GO) analyses. The results revealed activation of GO terms related to the mitochondria matrix [NES = 1.92, *p*.adjust = 0.007], the collagen-containing extracellular matrix [NES = 2.16, *p*.adjust = 0.029], and the extracellular matrix [NES = 2.14, *p*.adjust = 0.029] (Fig. 2D). In contrast, the suppressed GO terms included postsynapse [NES = −2.03, *p*.adjust = 0.001], asymmetric synapse [NES = −2.08, *p*.adjust = 0.035], postsynaptic density [NES = −2.05, *p*.adjust 0.039], postsynaptic specialization [NES = −2.07, *p*.adjust = 0.029], and neuron to neuron synapse [NES = - 2.22, *p*.adjust = 0.005] (Fig. 2D). Additionally, the actin cytoskeleton term was also suppressed [NES = −1.91, *p*.adjust = 0.034]. Further studies to elucidate the specific contributions of these pathways to AD pathogenesis will improve our understanding of the underlying pathophysiology of AD.

### Characterization of prolonged low-dose treatment in Aβ42A-treated iNs

The progression of AD is characterized by the increasing accumulation of misfolded Aβ peptides, which starts at low concentrations [29]. We treated iNs with Aβ42A at a low dose (0.5 µM) for a period of 21 days. This prolonged low-dose treatment significantly reduced the expression of neuronal markers, including MAP2, NEFH, and NEFM, while it upregulated the expression of AD-associated genes, such as APP, APOE, ADAM10, BACE1, and CLU (Fig. 3A & B). Differential protein abundance analysis further revealed more genes potentially involved in Aβ42A-induced neurotoxic processes, including the increased SFRP1, SULF2, SPOCK2, MDK, PTN, and SCUBE1, et al, as well as the decreased ARL6P4, SRRM2, TBC1D10B, SRRM2, PNN, and NOLC1, et al. (Fig. 3C). Notably, these molecular changes reproduced the observations of our previous study in which iNs were treated with a large dose of Aβ42A (5 µM), reinforcing the notion that Aβ42A induces specific cellular responses.

**Fig. 3.**
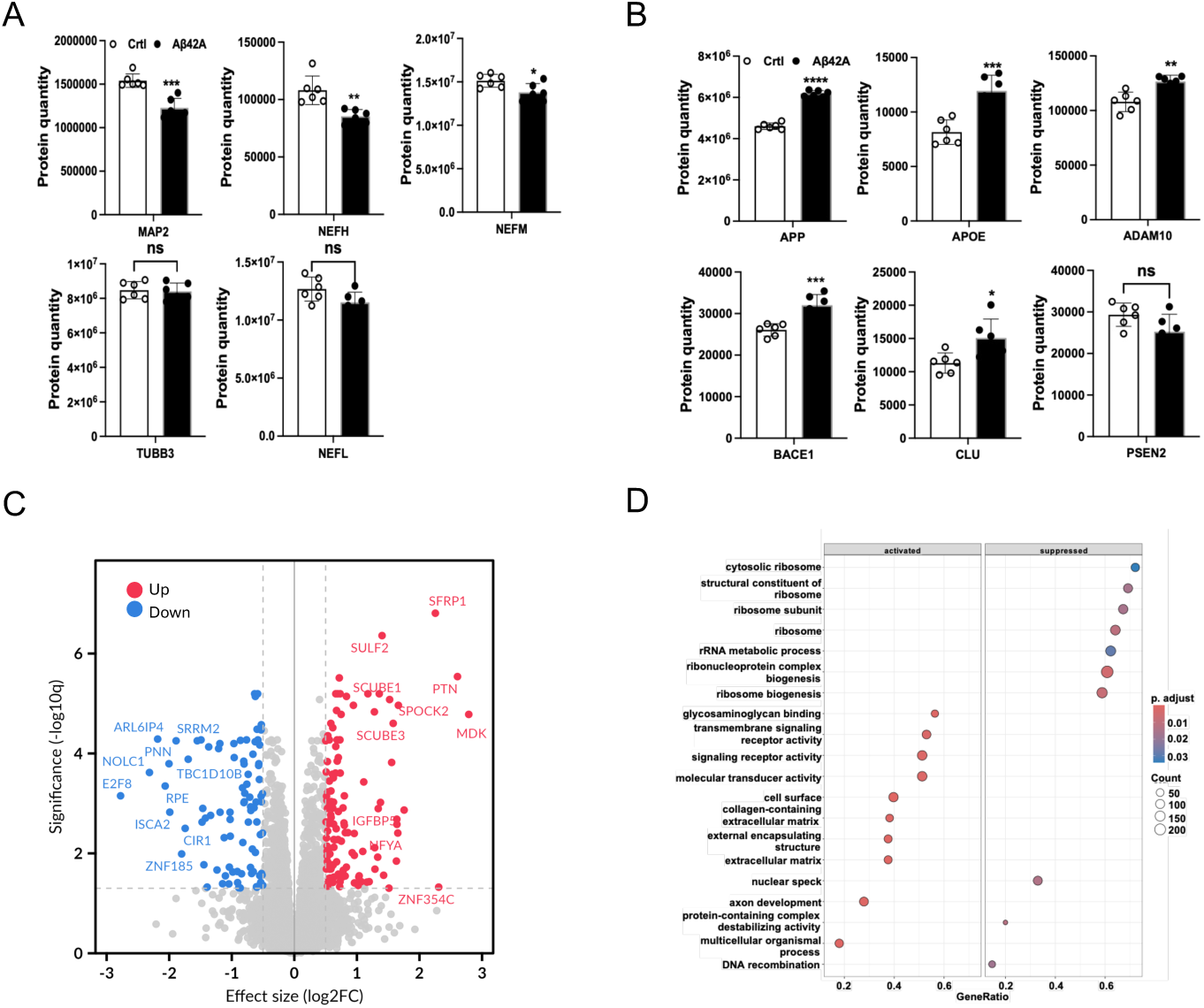
Prolonged low-dose treatment induced AD-related pathological alterations in KOLF2.1J iNs. A. Aβ42A regulated the expression of AD-related genes and cellular pathways. Day 7 iNs were treated with 0.5 µM Aβ42A for 21 days. The iNs were collected for LC-MS assay and proteomics analysis. Aβ42A treatment significantly downregulated the expression of multiple mature neuron markers, including MAP2, NEFH, and NEFM. B. Aβ42A treatment significantly upregulated several AD-associated molecules, including APP, APOE, ADAM10, BACE1, and CLU. C. Volcano plot of differential protein abundance analysis indicating the upregulated and downregulated proteins. D. Thematic classification of GO terms modulated by Aβ42A treatment.

To investigate the cellular pathways involved in iNs treated with a prolonged low dose of A42βA, we conducted GO analyses. The results revealed activated GO terms, including collagen-containing extracellular matrix [NES = 2.43, *p*.adjust = 0.0003], external encapsulating structure [NES = 2.42, *p*.adjust = 0.00002], and extracellular matrix [NES = 2.42, *p*.adjust = 0.00002] (Fig. 3D). Suppressed GO terms were associated with ribosomal functions and RNA processing, including cytosolic ribosome [NES = −2.24, *p*.adjust = 0.034], structural constituents of ribosome [NES = −2.10, *p*.adjust = 0.016], ribosomal subunit [NES = −2.08, *p*.adjust = 0.016], ribosome [NES = −2.06, *p*.adjust = 0.011], rRNA metabolic process [NES = −1.92, *p*.adjust = 0.029], ribonucleoprotein complex biogenesis [NES = −1.87, *p*.adjust = 0.005], and ribosome biogenesis [NES = - 1.98, *p*.adjust = 0.007] (Fig. 3D). Further studies are warranted to reveal the specific contributions of these pathways to AD progression in neurons.

### Aβ42A-induced spatially cellular responses in neuritic fractions

Neurons are highly polarized cells whose axonal and dendritic processes are collectively called neurites. However, their individual responses to Aβ aggregates remain unclear. To investigate the effects of Aβ aggregates on soma and neurites, we cultured iNs in Boyden chambers. After Aβ42A treatment, the NFs were collected first from the bottom of the Boyden chambers, while the remaining SEFs were harvested from the top of the Boyden chambers (Fig. 4A). The absence of nuclear proteins, such as NeuN, in NFs confirmed the efficient segregation of NFs and SEFs (Fig. 4B). Differential protein abundance analysis further demonstrates distinct proteomic changes between NFs and SEFs after Aβ42A treatment (Fig. 4C & D). The NFs of VC-treated iNs presented enrichment of many synaptic proteins, such as SYN1-3, SYT1-7, SYT9, and SYP (Fig. 4C), while Aβ42A treatment diminished those upregulated proteins in NFs (Fig. 4D). Moreover, more proteins were altered after Aβ42A treatment (Fig. 4C & D). The specific functions of these proteins in NFs are worthy of further investigation.

**Fig. 4.**
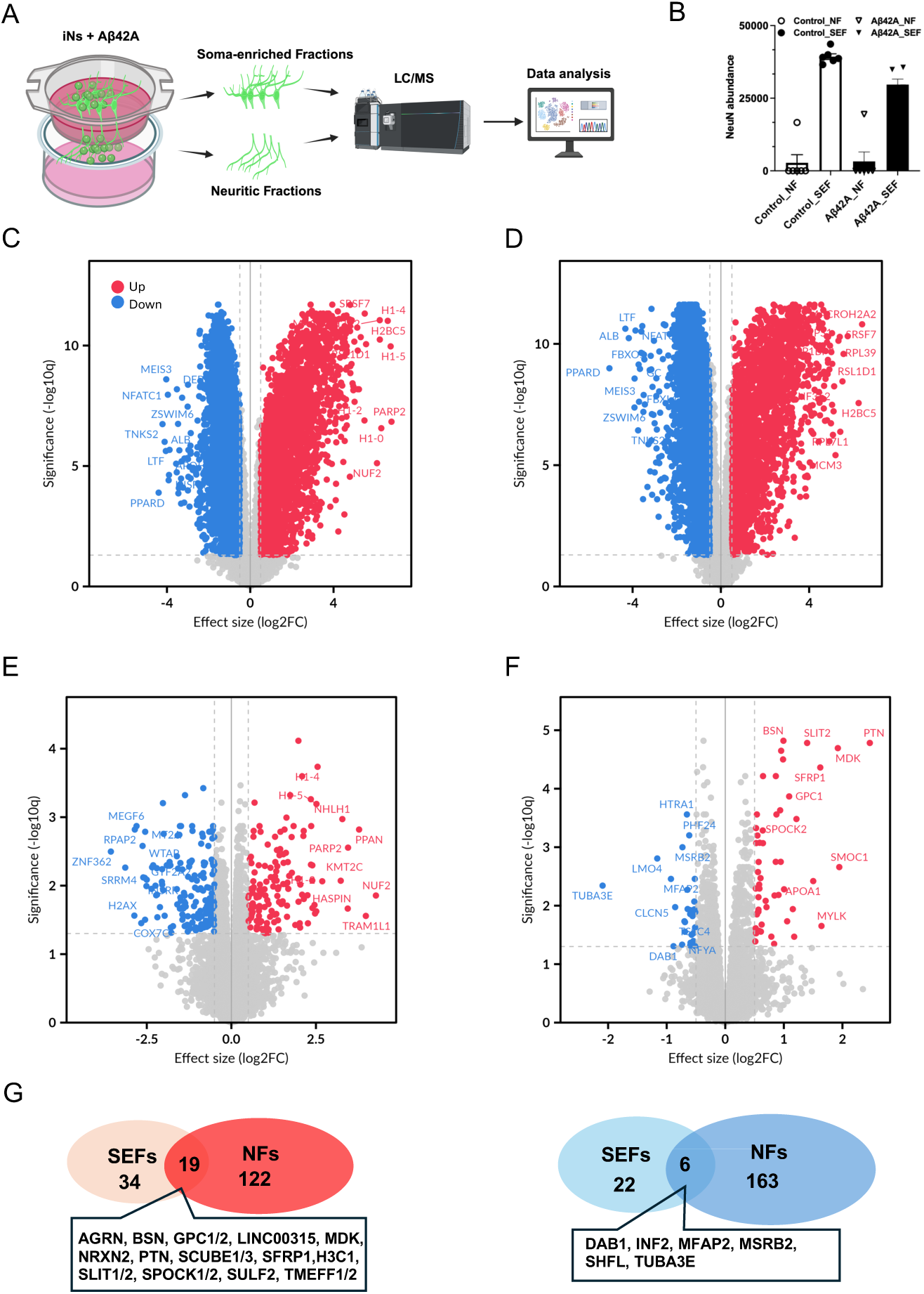
Aβ42A treatment impacted the NFs and SEFs of iNs in a spatially dependent manner. A. Scheme of the experiment design. Day 14 iNs were treated with 2.5 µM Aβ42A for 7 days. The NFs were collected first by scraping neurites from the bottom of the Boyden chamber inserts, and then the SEFs were harvested from the top of the inserts. The NFs and SEFs were processed for LC-MS assay and proteomics analysis. B. The abundance of NeuN in NFs and SEFs. C. Volcano plots comparing NFs and SEFs in VC-treated samples. D. Volcano plots comparing NFs and SEFs in Aβ42A-treated samples. E. Volcano plots of differential expression proteins in NFs, comparing VC-treated samples and Aβ42A-treated samples. F. Volcano plots of differential expression proteins in SEFs, comparing VC-treated samples and Aβ42A-treated samples. G. Venn diagram showing the overlap of upregulated proteins in NFs and SEFs. H. Venn diagram showing the overlap of downregulated proteins in NFs and SEFs.

Additional differential abundance analyses focus on the significant changes in proteins in NFs and SEFs separately. It revealed 141 proteins significantly upregulated in NFs and 53 in SEFs respectively and 19 proteins shared between both fractions (Fig. 4G). Several of these genes, including AGRN, BSN, MDK, NRXN2, PTN, SFRP1, and TMEFF2, have been implicated and studied in neurodegenerative disease [21, 30–34]. Moreover, 169 proteins were significantly downregulated in NFs and 28 were significantly downregulated in SEFs, with only 6 overlapping proteins (Fig. 4H). These three proteins, DAB1, MSRB2, and TUBA3E, play roles in neurodevelopment, the oxidative stress response, and microtubule dynamics, implicating that they could be relevant to the pathology of neurodegenerative conditions [35–37]. Our findings suggest that Aβ42A spatially regulates neuron responses, impacting molecular components commonly modulated in neurites and soma, as well as distinct proteins that are differentially regulated between the two regions. The further investigation of these proteins may reveal new targets for the rescue of functions of neurons affected by Aβ deposition.

### Correlation of translational manifestations with the Aβ42A-iNs system

The complex pathophysiology of AD hinders the identification of effective therapeutic targets [1, 22]. The proteomic datasets generated from our study may offer a basis for the discovery of therapeutic targets. We compared and analyzed four datasets from our proteomic profiles, including the large concentration, prolonged low-dose, NFs, and SEFs datasets. We found that 13 proteins (AGRN, GPC1, GPC2, MDK, NRXN2, PTN, SCUBE1, SCUBE3, SFRP1, SPOCK1, SPOCK2, TMEFF1, TMEFF2) were significantly upregulated across all four datasets after multiple test correction (Fig. 5A). Among these altered proteins, AGRN and SFRP1 have already been reported to be associated with AD pathogenesis [32, 38].

**Fig. 5.**
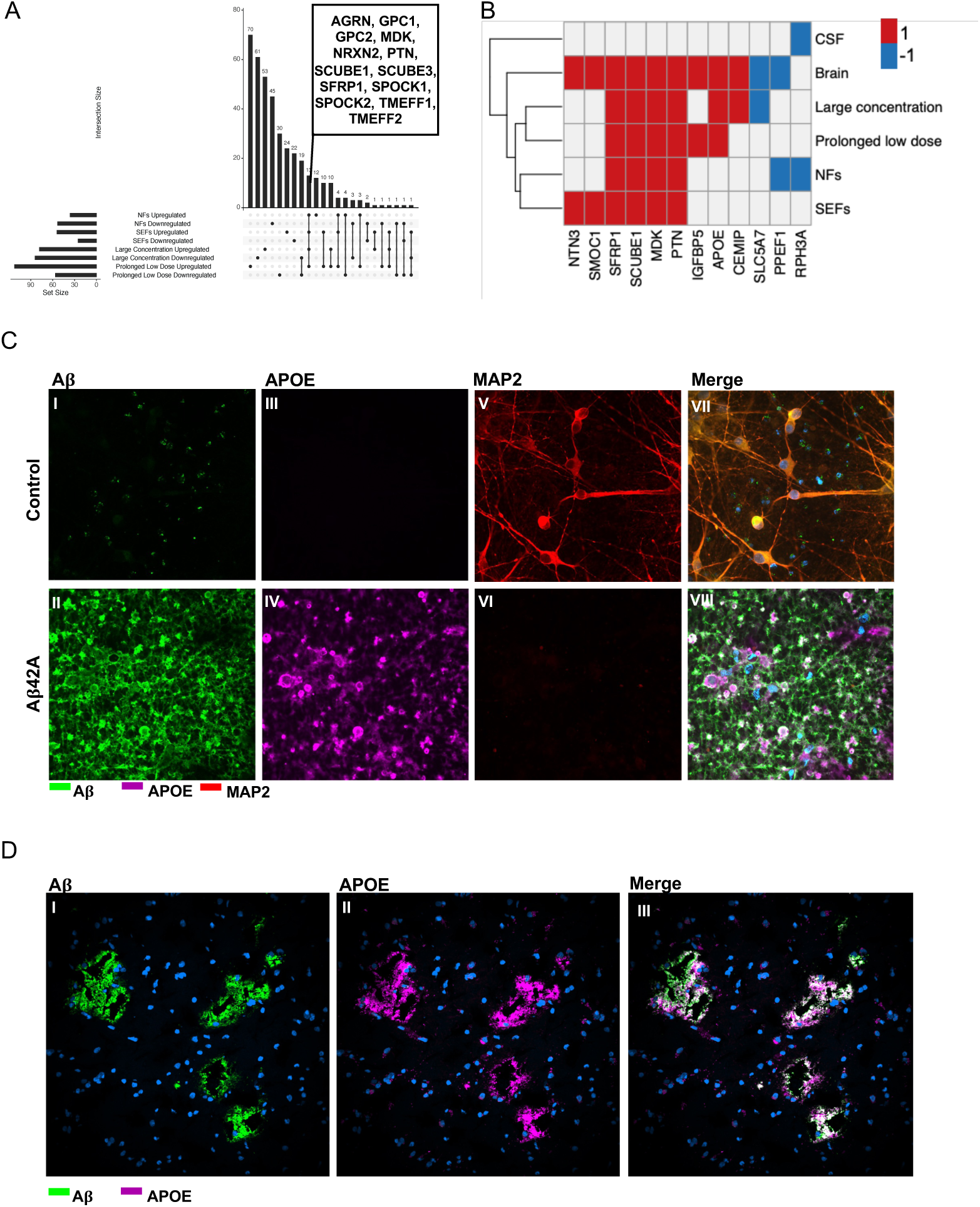
Replication of translational manifestations with the Aβ42A-iNs system. A. Upset plot showing the overlap of differentially expressed proteins among the four datasets. B. Heatmap showing commonly altered proteins between Aβ42A-treated iNs and human AD samples. C. Aβ and APOE staining in iNs. KOLF2.1J iNs were treated with Aβ42A and then fixed and stained with anti-Aβ and anti-APOE antibodies following fluorescent-conjugated secondary antibodies. The images were acquired using a Nikon confocal microscope. D. Aβ and APOE staining in human AD brain tissue. Frozen human AD brain tissues were fixed and stained with anti-Aβ and anti-APOE antibodies following fluorescent-conjugated secondary antibodies. The images were acquired using a Nikon confocal microscope.

To explore the translational relevance of our Aβ42A-iNs system, we compared our identified proteins with findings from postmortem Alzheimer’s disease brain tissues, which is a meta-analysis of seven deep proteomics datasets [18, 39]. The comparison identified 11 proteins (NTN3, SMOC1, SFRP1, SCUBE1, MDK, PTN, IGFBP5, APOE, CEMIP, SLC5A7, PPEF1) that were particularly dysregulated in both our findings (9568 proteins identified) and AD brain tissues (12017 proteins identified), and one protein (RPH3A) in our study (9568 proteins identified) and AD CSF (5939 proteins identified) (Fig. 5B). This overlap between our *in vitro* system and human samples provides validation that the Aβ42A-iNs system recapitulates intriguing molecular aspects of the neuronal pathophysiology of AD, supporting its utility as a relevant cellular platform for studying AD mechanisms and potential therapeutic interventions. For example, PTN, MDK, SCUBE1, and SFRP1 were consistently upregulated across all four of our datasets and in AD brain tissues, highlighting their uncovered importance in AD pathogenesis. APOE was another factor that was upregulated in two of our datasets and in AD brain tissues. The upregulation of IGFBP5 in the prolonged low-dose dataset, as well as the upregulation of CEMIP and the downregulation of SLC5A7 in the dataset from the large dose treatment, implied the specific regulation of AD pathogenesis by the dosage and exposure time of Aβ aggregates. Additionally, the upregulation of NTN3 and SMOC1 in SEFs, along with the downregulation of PPEF1 in NFs, suggested the spatial regulation of AD pathogenesis. Moreover, the significant downregulation of RPH3A in NFs and in AD CSF highlighted its potential as a diagnostic target. Overall, the alignment of our findings with those of clinical samples supports the validity and reliability of our Aβ42A-iNs system as a platform for mechanistic research and drug discovery.

To further validate the consensus of our findings in our system with the data of clinical samples. We investigated the APOE expression, as APOE is one of the strongest risk factors for AD, in both our cellular system and human brain tissues. Aβ42A treatment induced Aβ deposits (Fig. 5C II). Consistent with our mass spectrometry data, confocal imaging showed that the APOE expression was upregulated in Aβ42A-treated iNs (Fig. 5C IV). Additionally, the colocalization of APOE and Aβ42 was observed in Aβ42A-treated iNs (Fig. 5C VIII). The colocalization of APOE and Aβ42 in our cellular system intrigued us to investigate their distribution in the brain tissue from AD patient. In AD brain sections, Aβ plaques were readily detected (Fig. 5D I). Notably, distinct APOE-positive clusters were also observed in the tissue (Fig. 5D II), and the APOE clusters largely colocalized with Aβ plaques (Fig. 5D III). Together, these cellular and histological findings manifested that our system faithfully recapitulates the key features of AD pathology and supports its application for AD pathogenesis and therapeutic target identification.

## Discussion

Aggregated Aβ is a key driver of neurotoxicity and a primary therapeutic target for AD [1, 40]. However, the absence of standardized Aβ preparation protocol and the lack of refined forms of Aβ aggregates, plus the gap between recombinant Aβ aggregates and misfolded found in post-mortem brain tissues [41], post the challenge to produce a reliable cellular model. We utilized synthetic Aβ (1-42) peptides to establish a protocol for Aβ aggregates preparation, which was referred to as Aβ42A. Aβ42A aggregated in our *in vitro* iNs culture system and accumulated extracellularly around iNs. Aβ42A consistently induced neurotoxicity under different conditions. We are further characterizing the Aβ42A-iNs system to advance its translational application.

The genetic background plays important roles in AD. The KOLF2.1J iPSC line has been widely used in neurodegenerative disease research, although its inherent heterozygous copy-number variants may render line-specific effects and responses [11, 42–44]. We used KOLF2.1J iNs to study the neurotoxicity induced by Aβ42A. Compared with VC-treated iNs, Aβ42A-treated iNs exhibited impaired neurite outgrowth and decreases in various neuronal markers. Intriguingly, iNs showed a reciprocal response following Aβ42A treatment, characterized by the upregulation of both BACE1 and ADAM10. Moreover, a major AD risk factor, APOE, and a neurodegenerative disease marker, CLU, were upregulated in Aβ42A-treated iNs. Furthermore, Aβ42A treatment induced the dysregulation of a list of proteins, including NTN3, SMOC1, SFRP1, SCUBE1, MDK, PTN, IGFBP5, APOE, CEMIP, SLC5A7, PPEF1. These findings highlight the recapitulation of the neuronal pathophysiology of AD and reveal new potential regulators in AD pathogenesis by our Aβ42A-iNs system.

Our findings indicate that Aβ aggregates accumulate around the extracellular neurites and soma. However, it remains elusive whether Aβ aggregates elicit distinct cellular responses in neurites and soma. To address this, we separated neurites from neurons to generate NFs and SEFs and conducted proteomics analyses. We identified 169 proteins significantly downregulated in Aβ42A-treated NFs and 28 proteins downregulated in SEFs, with only 6 proteins shared between NFs and SEFs. We also observed a total of 141 proteins upregulated in Aβ42A-treated NFs and 53 proteins increased in SEFs, with 19 proteins overlapping between the two fractions. Our data demonstrated that Aβ42A spatially modulated cellular responses in neurites and soma. While a small set of these modulated proteins are reported to be related to AD pathogenesis, the remains have yet to be investigated for their roles in AD. Further studies to validate the roles of these molecules in AD pathophysiology may promote their potential as new therapeutic targets.

The human brain is composed of heterogeneous populations of cells [45, 46]. The complex relationship between neurons and nonneuronal cells increases the challenge of dissecting the contribution of a specific cellular population to AD pathogenesis [47, 48]. The iPSC-derived neuron system provides an opportunity to elucidate the mechanism of AD pathogenesis at the level of an enriched neuronal subtype. Our Aβ42A-iNs system, in which the majority consists of only neuron population, has replicated various aspects of neuronal pathophysiology of AD. To assess its translational relevance, we analyzed our findings in the context of clinical samples. We identified 12 proteins that showed significant changes in our observations, which also exhibited similar alterations in AD brain tissues or CSF. Among those proteins, APOE has long been recognized as a major risk factor for AD. We found the upregulation of APOE in our Aβ42A-iNs system with MS data and confocal images. Furthermore, we identified the colocalization of APOE and Aβ42A in both our culture system and in the AD brain tissue. Additionally, other proteins changed in Aβ42A-treated iNs, such as SFRP1, MDK, and PTN, have been reportedly involved in AD pathogenesis [21, 27, 32, 49]. Altogether, this consensus of our findings with clinical observations not only strengthens the validity of our Aβ42A-iNs system for mechanism studies but also suggests its potential for therapeutic discovery.

## Conclusion

We have established an AD cellular platform using iPSC-derived neurons and Aβ aggregates. This platform reproduces certain aspects of AD pathophysiology at the cellular level and aligns with clinical observations. It offers a valuable platform to bridge the gap in understanding how Aβ aggregates induce pathology in human neurons. Moreover, its potential for the discovery of therapeutic targets warrants further investigation.

## List of abbreviations

AD: Alzheimer’s Disease
iPSC: induced pluripotent stem cell
Aβ: amyloid β
iNs: induced pluripotent stem cell-derived neurons
Aβ42A: AggreSure amyloid β (1-42)
CSF: cerebrospinal fluid
NGN2: neurogenin-2
PLO: poly-L-ornithine
NMM: neuronal maturation media
PBS: phosphate-buffered saline
ACN: acetonitrile
LC/MS: liquid chromatography and mass spectrometry
NFs: neuritic fractions
SEFs: soma-enriched fractions
DIA: data-independent acquisition
FDR: false discovery rate
PSMs: peptide-spectrum matches
DEPs: differentially expressed proteins
GO: gene ontology
NES: normalized enrichment score

## Ethics declaration

Not applicable

## Consent to Participate

Not applicable

## Consent to Publish

Not applicable

## Disclamation declaration

This research was supported in part by the Intramural Research Program of the National Institutes of Health (NIH). The contributions of the NIH authors are considered Works of the United States Government. The findings and conclusions presented in this paper are those of the authors and do not necessarily reflect the views of the NIH or the U.S. Department of Health and Human Services.

## Data availability

The MS raw files and the processed protein abundance data supporting the conclusions of this article are available in the ProteomeXchange Consortium via the PRIDE partner repository, project PXD070927.

## Acknowledgments

This research was in part supported by the Intramural Research Program of the National Institute on Aging, National Institutes of Health, Department of Health and Human Services (Grant No. ZIAAG000534 to MRC).

The schematic diagram of this paper was generated with BioRender software (https://biorender.com/).

## Contributions

B.J. and Y.A.Q. conceived the idea, designed the experiments, analyzed the data, and wrote the manuscript; B.J., I. K., E.L., M.S. and V.H.R. conducted the experiments; Z.Y., J.E. I.K, E.L, V.H.R, and Y.H. analyzed the data and wrote a partial manuscript; Y.A.Q., B.J. E.L. and V.H.R. supervised the work; C.A.W., M.A.N., D.T., S.F., P.N., A.B.S., K.V.K., L.F., M.E.W., and M.R.C. revised the manuscript. All the authors have read, edited, and approved the final manuscript.

